# The tricyclic antidepressant cyproheptadine rescues egg-laying defect of *grk*-2 mutants

**DOI:** 10.1101/2025.05.09.653175

**Authors:** Jianjun Wang

## Abstract

Serotonin regulates egg-laying by activating vulva muscle contraction. Its oxidization metabolite 5-HIAA inhibits vulva muscle contraction. The loss-of-function mutants of *grk-2* have strong egg-laying defects because they have serotonin depletion and high level of 5-HIAA. The experiment results show that 5-HIAA’s function depends on 5-HT1/HT2 receptor SER-1. The egg-laying inhibition can be completely reverted by loss-of-function *goa-1*, the Gαo homolog in *C. elegans*. Serotonin and 5-HIAA competes for SER-1 binding and they activate different Gα protein complexes. The tricyclic antidepression drug cyproheptadine can block 5-HIAA function. The experiment results propose a plausible mechanism of cyproheptadine function. On the postsynaptic membrane, SER-1 binding with cyproheptadine can only recruit serotonin, thus, it biased activates Gαq instead of Gαo coupled signaling.

## Introduction

G protein-coupled receptor signaling controls many behaviors in eukaryotic organisms. Ligand binding to GPCR induces conformational changes that drive its interaction with heterotrimeric G proteins which promote the activation of downstream target proteins (1). GPCR kinases (GRKs) and arrestins downregulate the signal by inducing the endocytosis of GPCR complex (2). In model animal, *Caenorhabditis elegans*, grk-2 homolog to mammalian GRK2 & GRK3 was shown to play important roles in chemosensory (3,4) and egg-laying behaviors (5). In loss-of-function *grk-2* mutants, many unlaid eggs are held in uterus. The defect of egg-laying is caused by the oxidation of serotonin and its metabolite 5-HIAA inhibits vulva muscle contraction. The knockout strain of 5-HT2 homolog *ser-1(ok345)* is completely unresponsive to exogenous serotonin (6,7). It turns out that *ser-1* and *goa-1*are responsible for 5-HIAA function.

Tricyclic antidepression drug cyproheptadine is used to relieve the syndrome of allergy because it is against histamine signaling (8). Recently, cyproheptadine is used to cure appetite syndromes (9). Psychological depression is closely related to serotonin and its metabolite 5-HIAA (10,11). In our previous research, *grk-2* mutant worms have extremely low levels of serotonin and high levels of 5-HIAA. The animals look like they are in a great depression. It is interesting to investigate the mechanism of the syndrome and if the drug cyproheptadine could cure the diseases.

## Results

### The *grk-2* mutants are unresponsive to exogenous 5-HIAA

Previous study suggests that *grk-2 (rt97)* allele is loss-of-function mutant. It is induced by a single amino acid change T354I. The mutation could decrease its kinase activity dramatically (3). The other mutant *grk-2 (gk268)* has been knocked out of the first three exons of *grk-2* gene. Both mutants show strong chemosensory defects. Our group finds that *grk-2* mutants have egg-laying defect also. The strongest defect could be observed at 40 hours after vulva induction under the temperature 24°C (Figure 1A). Both mutated strains retain about 35 unlaid eggs in uterus. Under the same condition, wild type worm retains about 14 eggs. When 1mM 5-HIAA, the metabolite of serotonin, is added to the worm plate, the egg number of wild type worm increases to 21. However, there is no significant change in *grk-2* mutants (Figure 1A). The gain-of-function of *egl-30* (Gαq) has a big increase and the loss-of-function *goa-1* (Gαo) does not response to 5-HIAA (Figure 1A).

**Figure 1.**
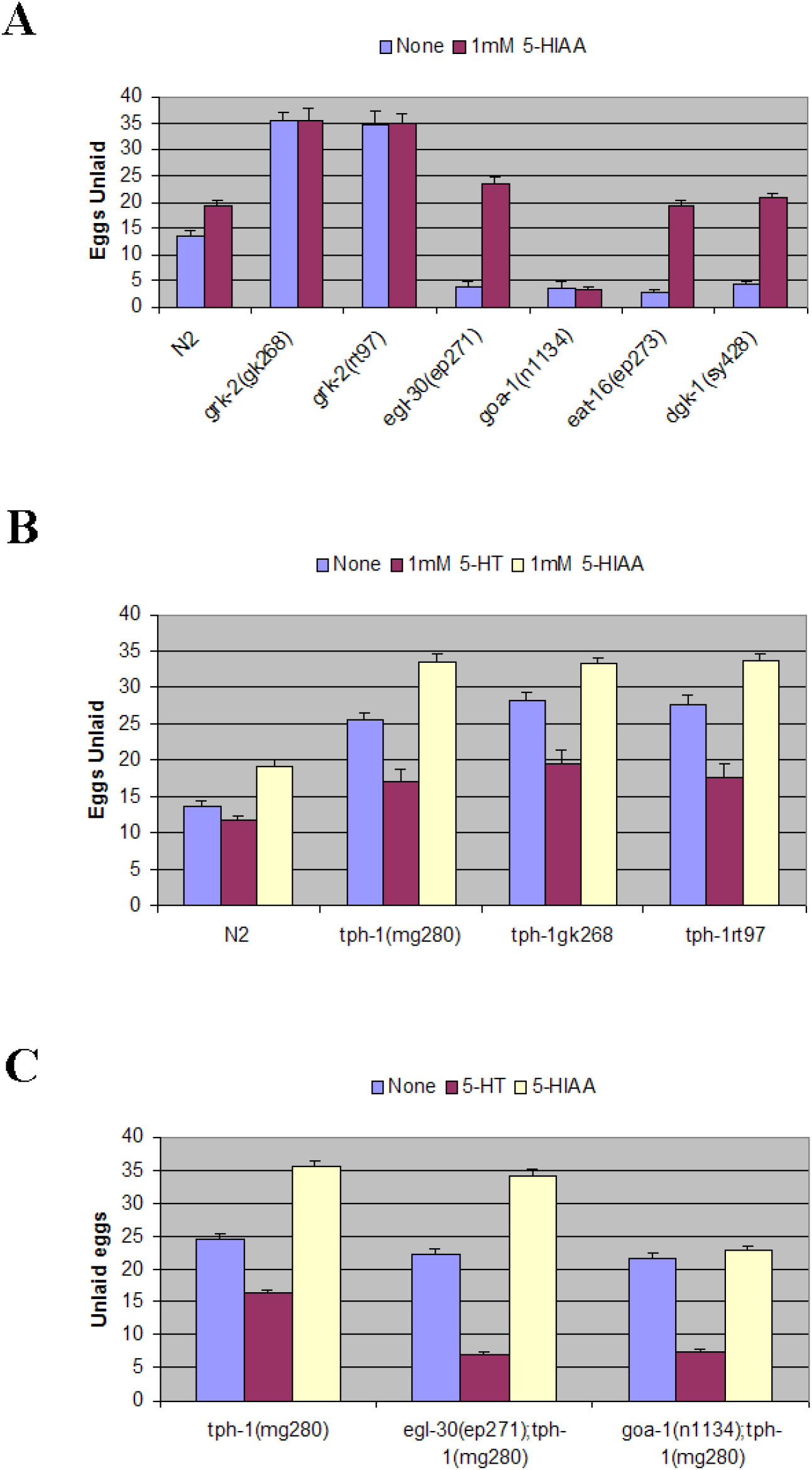
5-HIAA induces egg-laying defect in *grk-2* mutants via *goa-1*. A) *grk-2* mutants are defective in egg-laying, and they do not respond to exogenous 5-HIAA. Gain-of-function of *egl-30* is more sensitive to exogenous 5-HIAA and the loss-of-function *goa-1* has no response. B) The mutation of *tph-1* which is defective in serotonin synthesis, proves 5-HIAA can induce egg-laying defects. C) *tph-1;egl-30* and *tph-1;goa-*1 double mutants have different responses to exogenous 5-HIAA.

To further confirm this result, we crossed *grk-2* mutants with *tph-1*(mg280) which is defective in serotonin synthesis (12). The strain *tph-1*(mg280) has weak egg-laying defects and it retains about 25 eggs (Figure 1B). When fed on the plate with 1mM serotonin, the syndrome is completely rescued. However, when treated with 1mM exogenous 5-HIAA, the egg-laying defects are much stronger (Figure 1B), the unlaid egg number increased to 34. The double mutants of *grk-2; tph-1* have similar patterns. It suggests that without endogenous 5-HIAA, loss-of-function *grk-2* does not make the egg-laying defects stronger than *tph-1* alone. The egg-laying defect in *grk-2* mutants are induced by endogenous 5-HIAA. Its concentration could be close or higher than 1mM.

### GOA-1 is required in 5-HIAA signaling

The serotonin metabolite 5-HIAA is not a waste. It turns out to be a ligand binding GPR35 (orphan G protein0coupled receptor) (13,14). Then, 5-HIAA/GPR35 complex activates Gi/o proteins to inhibit inflammation. In worm egg-laying behavior, EGL-30(Gαq) and GOA-1 (Gαo) play important role in the neuron/muscle signaling (15,16,17,18). In figure 1A, *egl-30(ep271)* has gain-of-function mutation of EGL-30. The mutant worm has egg-laying constitutive phenotype, retaining less eggs than the wild type. When treated by 1mM 5-HIAA, its response is a little stronger than the wild type. The data from the loss-of-function Gαo/ *goa-1(n1134)* suggests that Gαo homolog protein GOA-1 is required for 5-HIAA function. 5-HIAA has completely no effect on egg-laying behavior of *goa-1(n1134)*. The double mutants of *egl-30(ep271); tph-1(mg280)* has enhanced response to exogenous serotonin, and its response to exogenous 5-HIAA is like *tph-1(mg280)* (Figure 1C). This suggests that EGL-30(Gαq) is important for serotonin signal, and it is not the target G protein of 5-HIAA. In contrast, the double mutant *goa-1(n1134); tph-1(mg280)* does not respond to exogenous 5-HIAA (Figure 1C), it suggests that GOA-1 is the downstream target G-protein of 5-HIAA signaling. The neurotransmitter 5-HIAA activates GOA-1, then turns on the Go/I coupled downstream signals including the inhibition of vulva muscle contraction.

### SER-1/5-HT GPCR plays dual roles in egg-laying

There are many G protein-coupled receptors in the worm genome (15). SER-1 is closely related to the 5-HT2 receptor which binds with serotonin and activates Gαq corelated signaling (6,7). The knockout strain *ser-1(ok345)* has a weak egg-laying defect and it does not respond to either exogenous serotonin or exogenous 5-HIAA (Figure 2A). The knockout strains of *ser-4* and *ser-7* have strong response to serotonin and 5-HIAA (data not shown). It suggests SER-1 is required for both serotonin and 5-HIAA signaling. The double mutant *grk-2(gk268);ser-1(ok345)* retains a few more eggs than *ser-1(ok345)*, and *grk-2(rt97);ser-1(ok345)* is completely same as *ser-1(ok345*). The egg-laying defects of the double mutants are much weaker than the *grk-2* mutants (Figure 2A).

**Figure 2.**
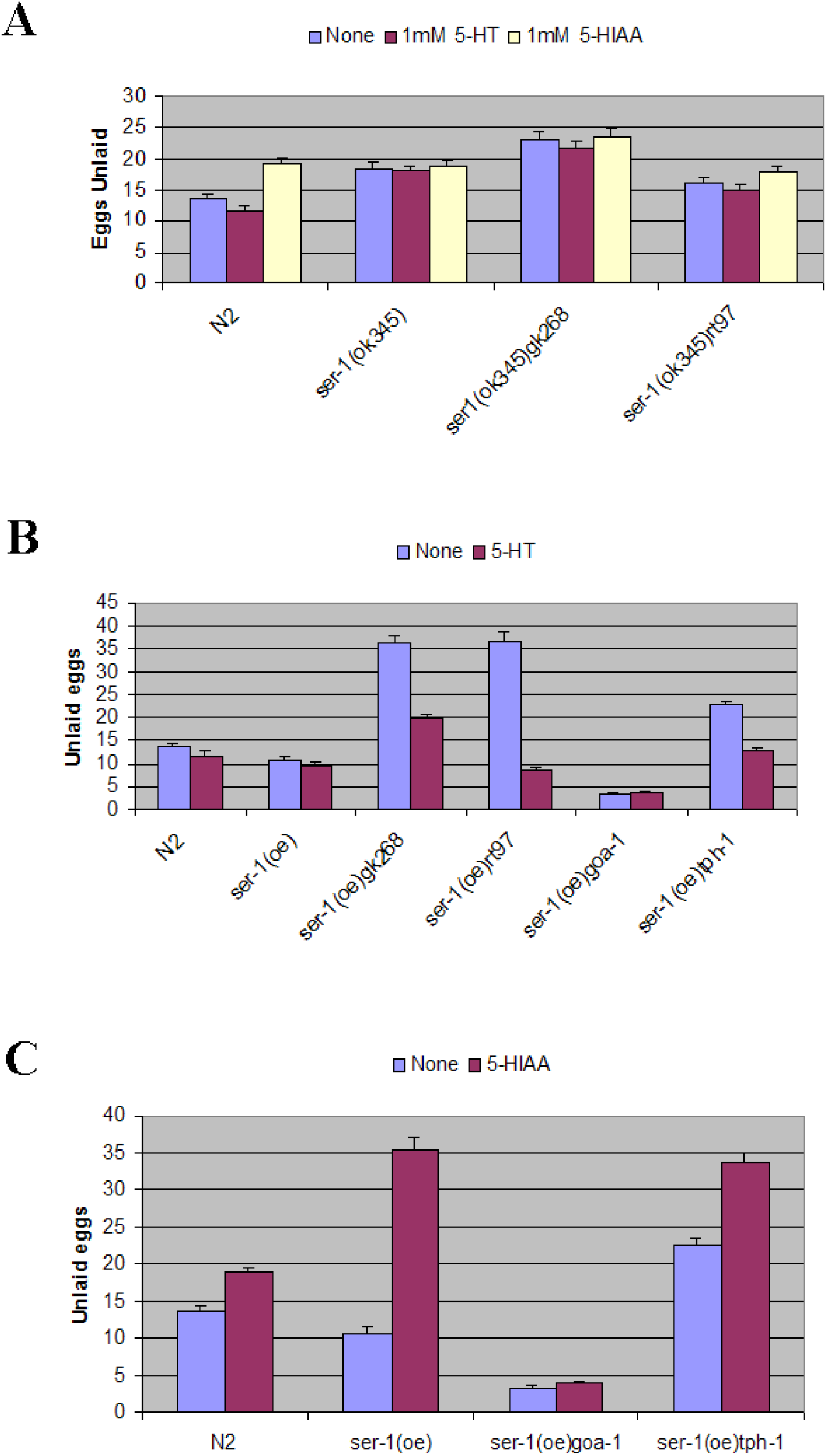
5-HIAA binds with SER-1 and activates GOA-1 complex. A) The knockout strain *ser-1(ok345)* mutant does not respond to endogenous and exogenous 5-HIAA. B) Overexpression of *ser-1* is more sensitive to exogenous serotonin. C) Overexpression of *ser-1* is more sensitive to exogenous 5-HIAA.

When *ser-1* is overexpressed using the extrachromosomal array in strain *ser-1(oe)*, the transgenic strain has a phenotype of egg-laying constitutive. The phenotype becomes stronger with 1mM exogenous serotonin. Without endogenous serotonin, in *tph-1(mg280);ser-1(oe)*, the egg-laying constitutive phenotype is completely suppressed. Interestingly, double mutants, *ser-1(oe);grk-2(gk268)* and *ser-1(oe);grk-2(rt97)* have egg-laying defects similar to *grk-2* mutants. The defects can be rescued by exogenous serotonin (Figure 2B).

More interestingly, when fed on the plate with 1mM 5-HIAA. The strain *ser-1(oe)* shows a strong defect of egg-laying like *grk-2* mutants which is much stronger than wild type worms (Figure 2C). It can be suppressed by loss-of-function of *goa-1* or *tph-1*(Figure 2C). It suggests that SER-1 could be the only GPCR in *C. elegans* whose ligand could be 5-HIAA. When 5-HIAA is released into the synaptic gap, SER-1 expressed on post-synaptic muscular membrane will bind 5-HIAA and the changed conformation activates Gαo which inhibits egg-laying.

### Cyproheptadine inhibits 5-HIAA function

Previously, cyproheptadine is reported to increase the worm lifespan under diet restriction condition (20). Our experiment also showed that this drug rescues the lifespan of *grk-2* mutants (data not shown). When 1uM cyproheptadine is added to the NGM plates, the wild type worms show strong egg-laying constitutive phenotype. A complete rescue of egg-laying defect of *grk-2(gk268)* mutant is observed when the worms are treated by cyproheptadine. Under the same condition, *grk-2(rt97)* shows egg-laying constitutive phenotype (Figure 3A). Because *egl-30(n686)* has a substitution mutation in Gαq protein it cannot be activated, and the mutant has a strong egg-laying defective phenotype. Interestingly, the defect can be suppressed by 1uM cyproheptadine (Figure 3A). Additional exogenous serotonin or 5-HIAA does not make any further change. The egg-laying constitute phenotype of *goa-1(n1134)* does not respond to cyproheptadine at all. It suggests GOA-1 is the target G protein for the cyproheptadine. More interestingly, cyproheptadine has no effect on *tph-1(mg280)* which cannot synthesize endogenous serotonin. With addition of 1mM serotonin, the mutant becomes egg-laying constitutive (Figure 3A). It suggests that the stimulation of egg-laying by cyproheptadine depends on the availability of serotonin. Addition of 1mM 5-HIAA does not change *tph-1(mg280)* at all. Comparingly, *tph-1(mg280)* fed with 1mM 5-HIAA alone which retains 34 eggs (Figure 1B). It means that the function of 5-HIAA is suppressed by cyproheptadine.

**Figure 3.**
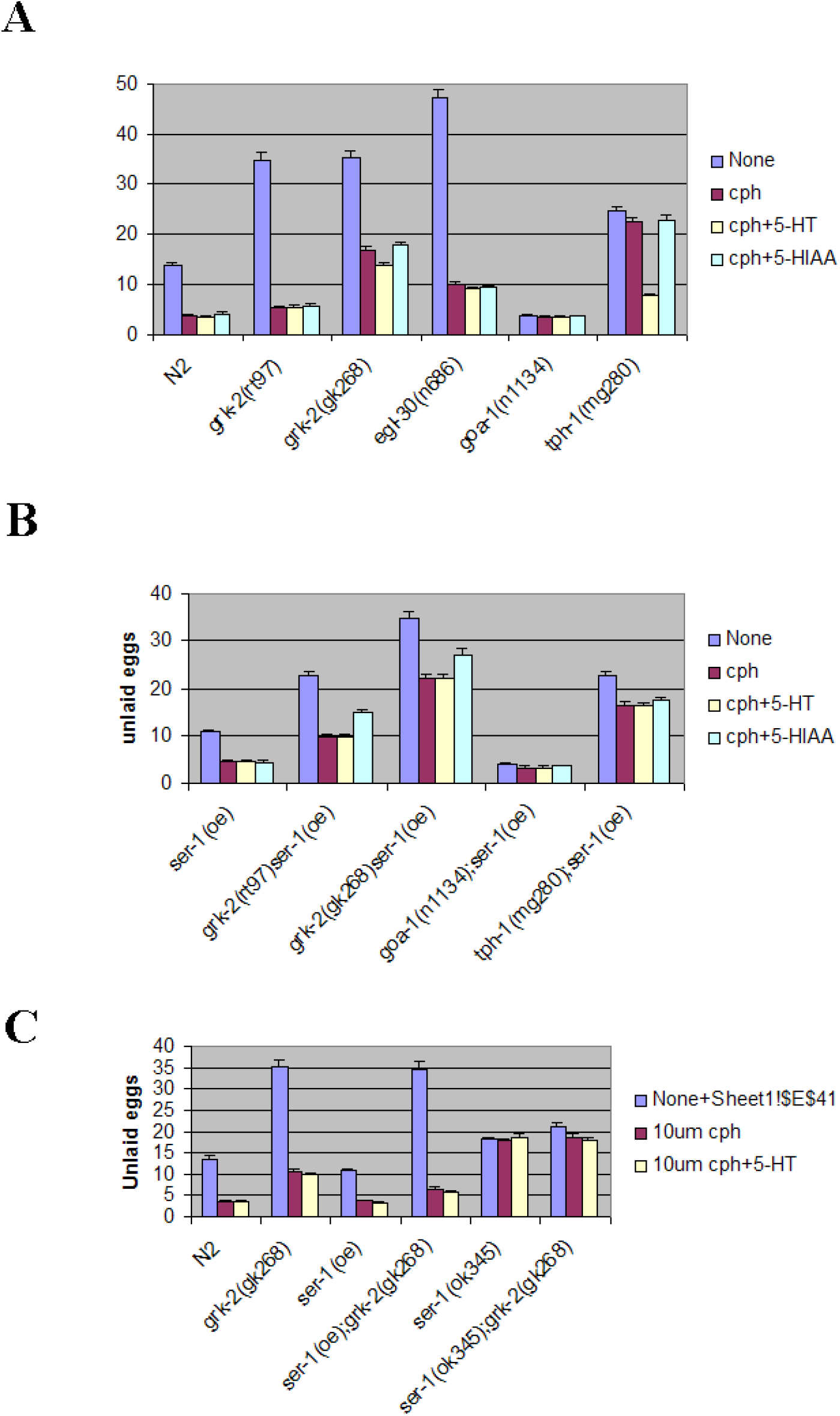
The tricyclic antidepressant cyproheptadine suppresses 5-HIAA function. A) 1uM cyproheptadine rescues egg-laying defects of *grk-2* mutants. The rescue depends on the availability of serotonin. B) When *ser-1* is overexpressed, 1uM cyproheptadine is not enough to suppress all 5-HIAA signal. C) The suppression needs the concentration of cyproheptadine increased to 10uM.

Overexpression of *ser-1* has egg-laying constitutive phenotype and it can be enhanced by adding 1uM cyproheptadine (Figure 3B). The double mutant *grk-2(gk268)ser-1(oe)* has a strong egg-laying defect like *grk-2(gk268)* which has saturated endogenous 5-HIAA and depleted serotonin level. Addition of 1uM cyproheptadine can only partially rescue this double mutant. The combination of 1uM cyproheptadine and 1mM serotonin has the same effect. With 1mM exogenous 5-HIAA, the rescue is even more trivial (Figure 3B). Under all conditions, *goa-1(n1134)* retains the same numbers of eggs. However, when the dosage of cyproheptadine increases to 10uM, its rescue of *grk-2(gk268)* becomes complete. The double mutant becomes egg-laying constitutive (Figure 3C). The knockout strain *ser-1(ok345)* does not respond to 10uM cyproheptadine. The drug cyproheptadine rescuing *grk-2* mutant phenotype depends on GPCR protein SER-1.

## Discussion

### 5-HIAA is responsible for muscle relaxation

Under normal condition, serotonin is synthesized in neurons including HSNs (6). With upstream stimulation, serotonin is released into neuron-muscle synapse and binds with post-synaptic cellular membrane anchored serotonin receptors including SER-1(7). The activated GPCR forwards the signaling to target G protein complex. Serotonin will be reuptake and oxidized into 5-HIAA by neurons. To prevent the damage from the over exhausting signaling, the competition between 5-HIAA and serotonin binding to the same GPCR is an effective method. When the endogenous 5-HIAA level is high, it suggests that the muscle could be exhausted, and it needs relaxation.

### 5-HIAA binds with SER-1 and activates GOA-1

In all experiments, 1mM serotonin or 1mM 5-HIAA is directly added into the NGM plates. The knockout strain *ser-1(ok345)* completely removes the GPCR *ser-1* which is a close homolog of both human 5-HT1 and 5-HT2 receptors. It is confirmed by protein sequence alignment (data not shown). Thus, it is not a surprise that *ser-1(ok345)* does not respond to any external compound including serotonin, 5-HIAA and cyproheptadine. The overexpression of *ser-1* has enhanced response to exogenous serotonin and 5-HIAA. Since 5-HIAA functions as an inhibitor of vulva muscle contraction and it depends on GOA-1. Thus, 5-HIAA binds on SER-1 and activates GOA-1 coupled signaling is the most reasonable hypothesis.

### Cyproheptadine inhibits 5-HIAA/GOA-1 coupled signaling

The compound cyproheptadine is a very effective drug to activate egg-laying. Its function depends on the availability of serotonin, either endogenous or exogenous. It suggests cyproheptadine regulates egg-laying by enhancing serotonin signaling. At same time, cyproheptadine suppresses 5-HIAA signaling. Since serotonin and 5-HIAA compete for SER-1 binding, it suggests that cyproheptadine can bind on SER-1. After binding with cyproheptadine, with a specific conformational change, SER-1 receptor is more sensitive to serotonin, and does not bind with 5-HIAA any longer. Cyproheptadine is an inverse agonist of 5-HIAA/SER-1/GOA-1 signaling.

In the knockout strain of *grk-2(gk268)*, the level of endogenous 5-HIAA could be extremely high. Its effect is even stronger than 1mM exogenous 5-HIAA. Treatment of 1uM cyproheptadine only relieves the egg-laying defect close to the wild-type level. When *ser-1* is overexpressed, 1uM cyproheptadine is not enough to block all receptors. Increasing the concentration of cyproheptadine ten times, the overall effect is similar to *grk-2(gk268)* alone. All data suggest that cyproheptadine is competitor of 5-HIAA for SER-1 binding.

## Materials and Methods

The compounds, serotonin and 5-HIAA are ordered from sigma-Aldrich. Cyproheptadine is from Fisher Scientific. Worm strains of individual mutants are ordered from CGC.

### Worm culture

All worms are cultured using nematode growth media (NGM) plates. For 1-liter NGM media includes 20g agar, 975g water, 5g peptone, 2.5 g tryptone, and 3g NaCl. For the plate with serotonin, 5-HIAA, or cyproheptadine, the compound is dissolved in di-water 1,000X stock solution and filtered by 0.22 um membrane. Add 1ml stock solution when NGM plate is cooled down to 55°C, after mixing well, pour the media into worm plates. All worms are grown under temperature at 24°C fed by E. Coli strain OP50.

### Worm crossing

To make double mutants or backcross the mutant strains, the male worms are generated by growing the L4 hermaphrodite worms under 33°C for 3 hours, then, pick 3 worms into new plates. Next generation will contain many male worms. For each cross, put three L4 hermaphrodite worms and three males on a plate with a small drop of E. Col at the center of the plate. After overnight incubation, each hermaphrodite will be transferred into individual plate. Next generation, successful crossed worms should contain 50% male worms. Pick three L4 hermaphrodite F1 generation worms and put them into three new plates, Monitor the growth of F2 generation worms, confirm the genotypes by DNA sequencing.

### Egg-laying assay

The worms are grown under low density, 2-3 hermaphrodite per plate at 24°C. After 48 hours, 12-15 L4 worms are transferred to a new plate. After another 24 hours, the adult worms are transferred into new plates again. After 16 hours, the egg retained in uterus is counted. The stage of each egg is determined under dissecting microscope.

